# Rapamycin Enhances CAR-T Control of HIV Replication and Reservoir Elimination *in vivo*

**DOI:** 10.1101/2024.10.31.621350

**Authors:** Wenli Mu, Shallu Tomer, Jeffrey Harding, Nandita Kedia, Valerie Rezek, Ethan Cook, Vaibahavi Patankar, Mayra Carrillo, Heather Martin, Hwee Ng, Li Wang, Matthew D. Marsden, Scott G. Kitchen, Anjie Zhen

## Abstract

Chimeric Antigen Receptor (CAR) T cell therapy has emerged as a powerful immune therapy for various diseases. Our studies in humanized mice and non-human primates (NHPs) demonstrate that hematopoietic stem cell (HSCs) modified with anti-HIV CAR leads to lifelong engraftment and supply of functional anti-viral CAR-T cells, leading to significantly reduced viral rebound after ART withdrawal. However, T cell exhaustion, driven by chronic immune activation, remains a major challenge for the continuous efficacy of CAR-T therapy, necessitating additional measures to achieve functional cure. We recently showed that *in vivo* treatment with low dose rapamycin reduced chronic inflammation and improved anti-HIV T cell function in HIV-infected humanized mice. Here, we report that rapamycin significantly improved CAR-T cell function both *in vitro* and *in vivo*. *In vitro* treatment with rapamycin improved CAR-T cell mitochondria respiration and cytotoxicity. *In vivo* treatment with low-dose rapamycin in HIV-infected, CAR-HSC treated mice reduced chronic inflammation, prevented exhaustion of CAR-T cells and improved CAR-T control of viral replication compared to CAR-HSCs treatment alone. RNAseq analysis of sorted CAR-T cells from humanized mice showed that rapamycin significantly modified the CAR-T cell transcriptome, including the downregulation of multiple check point inhibitors and the upregulation of key genes related to cell survival. We also observed significantly delayed viral rebound after ART withdrawal and diminished HIV reservoir in mice that were treated with rapamycin and CAR-HSCs as compared to CAR-HSCs treatment alone. Taken together, our data indicate that HSCs-based anti-HIV CAR-T combined with rapamycin treatment is a promising approach for treating persistent inflammation and improving immune control of HIV replication.

## INTRODUCTION

Engineering T cells with anti-HIV chimeric antigen receptors (CAR) has emerged as a promising gene therapy strategy to control HIV infection. HIV-specific CD8+ cytotoxic T lymphocytes (CTLs) are essential in suppressing HIV replication and eliminating HIV infected cells (1, 2). However, due to immune evasion by HIV (3) and development of dysfunctional HIV-specific T cells, natural CTLs are incapable of complete control of HIV replication in the absence of combination anti-retroviral therapy (ART) (4) and cannot eliminate reservoirs with “kick-and-kill” HIV cure strategies (5). A promising approach to overcome these barriers is through chimeric antigen receptor (CAR) engineered T cell therapy (6). T cells engineered with CD4-based CARs, which utilize CD4 extracellular domains to recognize HIV-1 Env, can effectively kill HIV infected cells and limit viral escape, as an escape from CD4 recognition would directly decrease viral fitness (7) (8). However, persistence, trafficking, and maintenance of function remain major challenges for peripheral CAR-T cell therapy (9). To overcome these issues, we showed that CAR modified hematopoietic stem cells (HSCs) are capable of lifelong engraftment and allow development of functional CAR-T cells *in vivo* (*10–12*). Our studies in humanized mice (10, 11) and non-human primates (NHPs) (12, 13) demonstrated the feasibility and efficacy of the CAR-HSC therapy and showed that CAR-HSCs transplanted animals have significantly reduced viral rebound after ART withdrawal. We have also made substantial improvements to this therapy by modifying the original CD4CAR construct to a second-generation CAR: termed D1D2CAR41BB. We showed that D1D2CAR41BB-HSC transplanted animals have improved CAR-T cell differentiation, better CAR-T cell persistence, and enhanced viral control (10). However, our lead CAR-HSC therapy still fell short of achieving complete viral remission in the absence of ART.

T cell exhaustion remains a major challenge for CAR T therapy for HIV-1 cure. Driven by chronic immune activation, T cell exhaustion remains one major barrier to achieving sustained immune surveillance for viral infection and cancer (14). Despite our recent successes in HSC-based CAR therapy, we found that HSCs-derived D1D2CAR41BB T cells are also subject to becoming exhausted. Similar CAR T cell exhaustion was also observed during peripheral anti-HIV CD4CAR T cell therapy in NHP models (15, 16). Although antibodies targeting ICBs (immune checkpoint blockades, such as PD-1 blockade) may restore CAR T cell function transiently (16, 17), ICB treatment can lead to serious side effects such as onset of type 1 diabetes, colitis, and other adverse effects (18, 19) that may be unacceptable to ART treated HIV+ individuals. Therefore, alternative strategies to safely reverse or prevent CAR-T cell exhaustion are critical for achieving long-term control of HIV replication.

Rapamycin, which inhibits mammalian Target of Rapamycin (mTOR) Complex 1, was first approved for use in anti-cancer therapies and transplant rejection prevention. In recent years, rapamycin has shown robust geroprotective ability, extending life spans in multiple model organisms, including the life span of genetically heterogeneous mice from multiple research groups (reviewed in (20)). In humans, rapamycin administration has been shown to reverse immunosenescence and boost response to seasonal flu vaccines (21, 22) and it is being studied in multiple clinical trials in healthy individuals and individuals with age-related diseases (23–25). Importantly, rapamycin can lead to metabolic reprogramming in T cells, shifting metabolism from glycolysis to oxidative phosphorylation (OXPHOS) and modulating lipid metabolism to enhance CD8 T cell memory formation (26, 27). While chronic treatment of humans with high doses of rapamycin or analogs is associated with deleterious metabolic effects, recent studies in humans have shown that a lower or intermittent dosing regimen of mTOR inhibitors in older adults is well tolerated and leads to improved immune function and reduced infection in the elderly (21, 22, 28). A recent study also demonstrated that rapamycin treatment during the beginning of chronic infection improves CD8 T cell memory formation and the efficacy of PD-1 targeted therapy (29). Moreover, our recent work showed that intermittent *in vivo* treatment of HIV+ humanized mice with rapamycin led to reductions in immune activation and improved endogenous anti-HIV T cell function, resulting in accelerated viral suppression during ART and reduced viral rebound after ART withdrawal (30). Here we show that low dose, intermittent rapamycin restores and improves anti-HIV CAR-T cell function during chronic HIV infection. We found that rapamycin treatment significantly remodeled the CAR-T cell transcriptome and improved mitochondria function, resulting in enhanced anti-viral activities of CAR-T cells. This led to delayed viral rebound after ART withdrawal and significantly improved viral control by CAR-T cells, suggesting potential therapeutic values of rapamycin in improving CAR-T cell function *in vivo*.

## RESULTS

### Rapamycin treatment normalized anti-HIV CAR-T mitochondria metabolism and improved CAR-T function *in vitro*

To examine if rapamycin modifies CAR T cell metabolism and restores exhausted anti-HIV-1 T cell functions *in vitro*, we investigated the effects of rapamycin on D1D2CAR41BB modified primary T cells that can recognize and kill HIV infected cells as described previously (10–12). CAR T cells were generated by activating primary PBMCs from HIV seronegative individuals and transducing them with a CAR expressing lentiviral vector. Afterward, CAR T cells were cultured *in vitro* for 2 weeks with IL-2 to stimulate T cell proliferation and induce T cell exhaustion. After culture, cells were treated with either rapamycin or vehicle control for 2 days followed by mitochondrial respiration analysis using a Seahorse assay. As shown in **Fig. 1A**, treatment with rapamycin significantly increased both basal (**Fig. 1B**) and maximal mitochondria respiration levels (**Fig. 1C**), highlighting its potential ability to rescue ATP-linked mitochondrial respiration in antiviral CAR-T cells. Rapamycin-treated CAR-T cells also showed a reduction in mitochondria reactive oxygen species (ROS) by MitoSOX staining (**Fig. 1D**). These results suggest that rapamycin can reduce ROS and enhance mitochondria functions in exhausted anti-HIV CAR T cells. To investigate if rapamycin has a restorative effect on anti-HIV-1 CAR T cell function, we tested rapamycin-treated CAR T cells’ ability to produce pro-inflammatory cytokines and cytotoxic capacity by co-incubating CAR T cells with control or target cells that express HIV envelope. As shown in **Fig. 1E** and **Fig.1F**, we observed significant increases in both IL-2 and IFNγ production and improved cytotoxic killing activity by CAR T cells treated with rapamycin. Taken together, these data strongly suggest that rapamycin treatment can remodel anti-HIV CAR-T cell metabolism and restore anti-viral T cell function *in vitro*.

**Figure 1.**
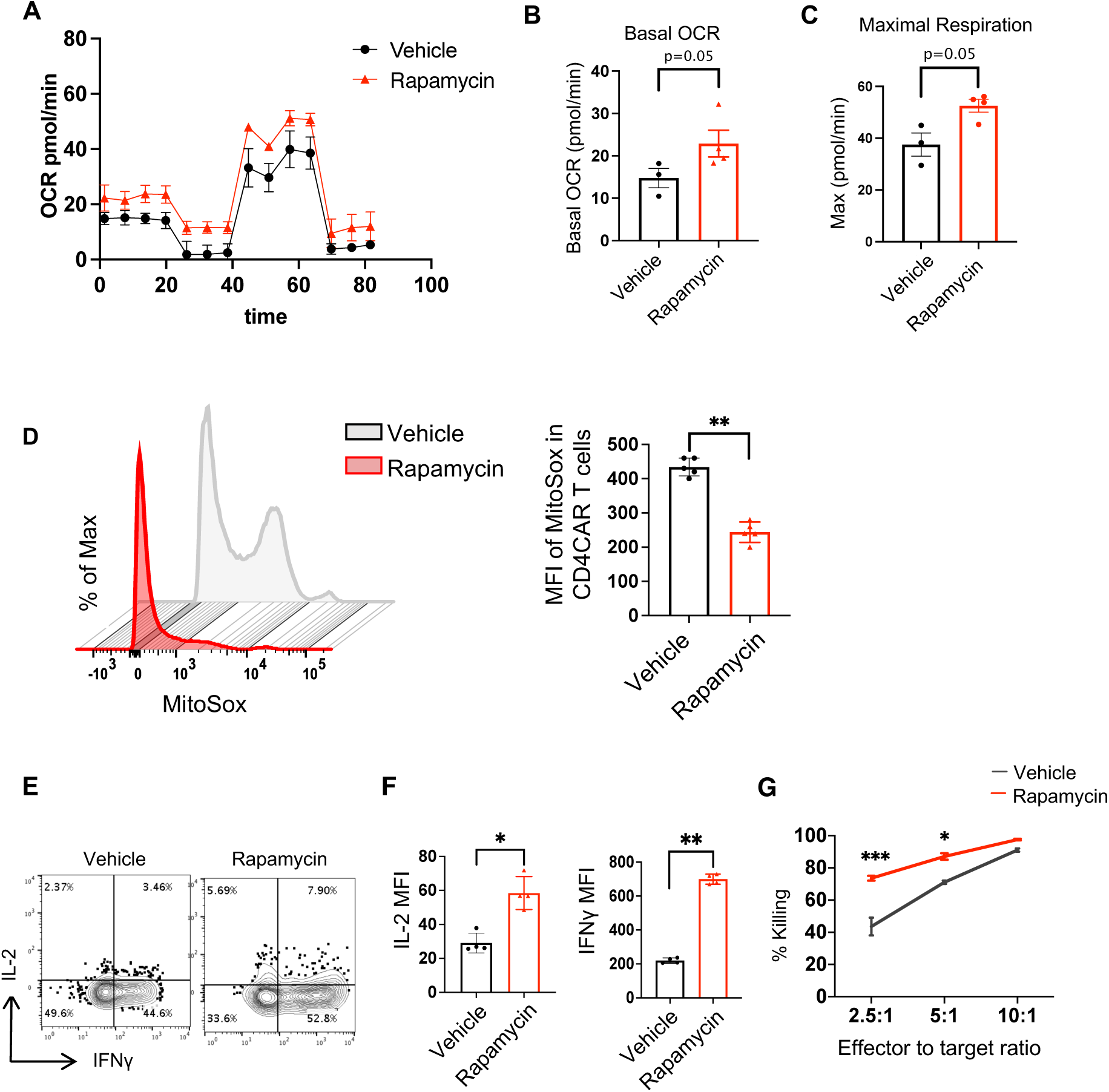
Treatment of anti-HIV CAR-T cells with rapamycin modified cellular metabolism *in vitro*. anti-HIV CD4CAR-T cells were produced by transducing activated primary PBMCs from healthy donors. Cells were then sorted to >90% CAR+ purity and expanded using 100IU/ml IL-2 for 2 weeks to promote exhaustion, followed by treatment with either DMSO, or 50pM of Rapamycin for 2 days. Afterwards, seahorse assay was performed on treated CAR T cells. **A**) The oxygen consumption rates (OCRs) under basal metabolic conditions and in responses to metabolic inhibitors. **B**) Basal OCR levels. **C**) Maximal respiratory levels. **D**) ROS were analyzed in CD4CAR-T cells labeled with MitoSOX after treatment as shown by flow cytometry and MFI summary of MitoSox. **E-F**) Cytokine assay. CAR-T cells were stimulated with HIV Env expressing (stimulated ACH2) cells overnight, followed by Golgiplug for 6 hours. Percentage of IFN-γ and IL-2 expression were measured by flow cytometry in CD4CAR-T cells. Representative flow plot and summary was shown in E and F. **G**) Killing assay. CAR-T cells were stimulated with HIV Env+ (stimulated ACH2) target cells or Env-(unstimulated ACH2) cells overnight at 2.5:1, 5:1 and 10:1 ratio. Specific killing activity is shown for vehicle treated and rapamycin treated CAR-T cells. All values are means ± SD of at least three independent experiments. Mann-Whitney test (unpaired); \**P* < 0.05, \*\**P* < 0.01, \*\*\**P* < 0.001.

### HIV induced HIV-specific CAR T exhaustion and dysfunction, while rapamycin treatment alleviated exhaustion and restored the viral suppression function of CAR T cells

Driven by chronic antigen stimulation and persistent immune activation, T cell exhaustion remains one major barrier to achieving sustained immune surveillance for chronic viral infection (31, 32). Similarly, CAR-T cell exhaustion is observed during peripheral anti-HIV CAR T cell therapy in both humanized mice (15) and NHP models (16), undermining the efficacy of CAR-T therapy. Despite our recent successes in inhibiting HIV replication in humanized mice and NHPs using HSCs-based CAR therapy (10–12) and improved CAR-T cell memory formation with a second generation CAR containing the 41BB domain (10), viral loads rebounded in all CAR-HSCs transplanted animals after ART removal. Therefore, it is crucial to investigate immune exhaustion of HSC-derived CAR T cells in chronic HIV infection *in vivo*. Hence, we constructed humanized bone marrow-liver-thymus (BLT) NSG mice with HSCs mock transduced or transduced with lentiviral vectors expressing D1D2CAR41BB, as described previously (10). After immune reconstitution, we infected BLT mice with a high dose of HIV-1_NFNSXL9_ (500ng of p24) to drive high viral loads and faster immune exhaustion (schematically shown in **Fig. 2A**). As expected, we observed a significant and gradual increase in the activation markers HLA-DR (**Fig. 2B**) and CD38 (**Fig. 2C**) on CAR T cells from peripheral blood after HIV infection (Representative flow gating strategy is shown in **SFigure1**). 3 weeks post infection, we also observed a significant increase in the exhaustion markers PD-1 and Tim-3 among CAR T cells (**Fig 2D-E)**. Consequently, while CAR mice demonstrated lower viral loads compared to mock 1 week after HIV infection, the viral suppression effects were lost by 3 weeks (**Fig 2F**), suggesting exhaustion/dysfunction of CAR T cells.

**Figure 2.**
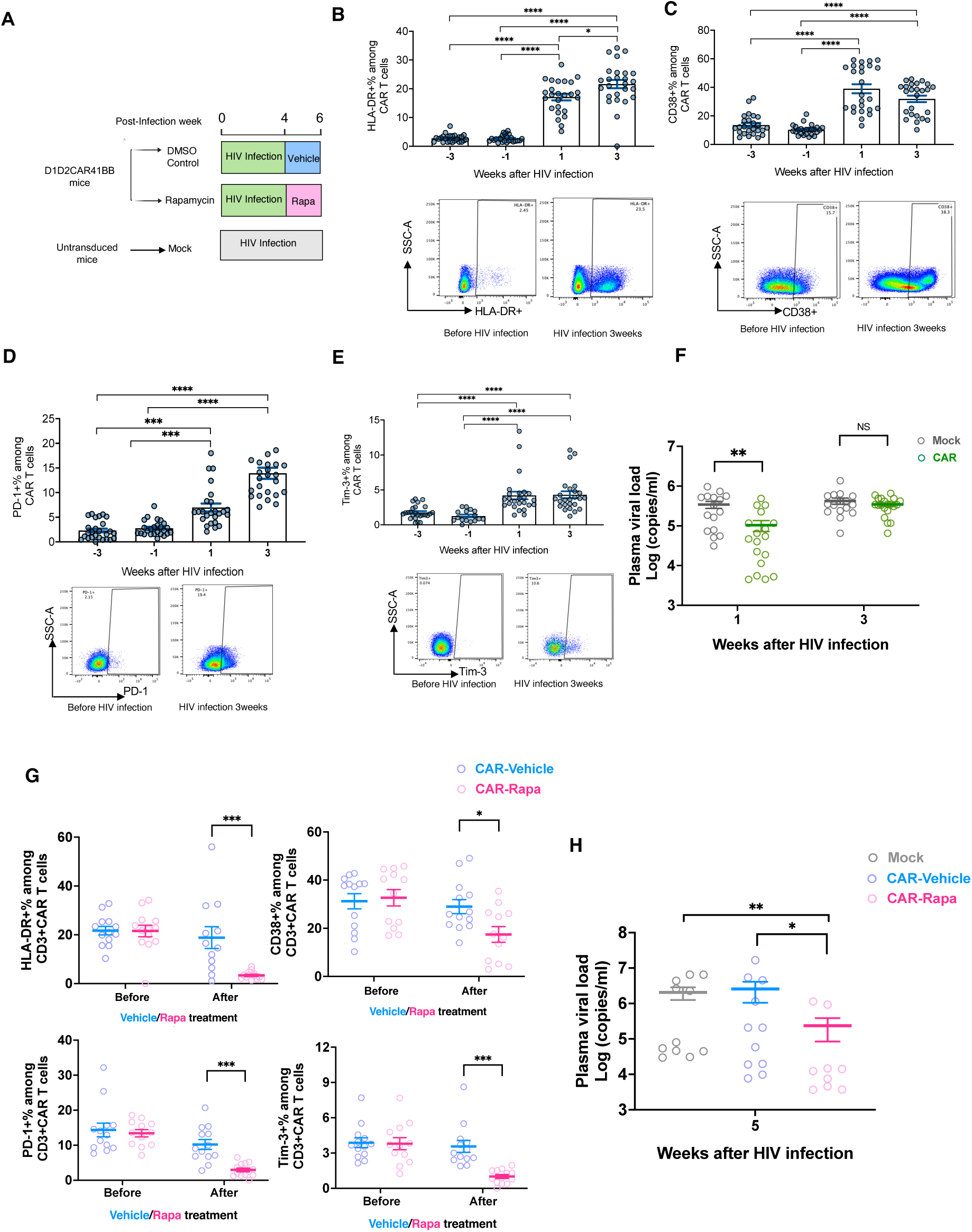
Chronic HIV infection leads to CAR-T cells exhaustion, while rapamycin treatment alleviates activation and exhaustion of CAR-T cells *in vivo*. **A**) Humanized NSG-BLT mice were constructed with either unmodified HSCs or HSCs modified with D1D2CAR41BB. After immune reconstitution, mice were infected with HIV-1_NFNSXL9_. Four weeks after infection, mice with CAR modified HSCs were treated with rapamycin or vehicle for 2 weeks (n=8-10). (**B-E**) Representative flow and average percentage of B) HLA-DR, C) CD38, D) PD-1, E) Tim-3 expression among CAR+CD3+ T cells before and after HIV infection as measured by flow cytometry (quantified by gating of percentage positive ± SEM). (**F**) Plasma HIV viral load from mock or anti-HIV CAR mice at 1 week and 3 weeks of infection. (**G**) Average percentage of PD-1, Tim3, HLA-DR and CD38 expression among blood CAR+CD3+ T cells before and after rapamycin treatment. (**H**) Plasma HIV RNA copies from mock mice or CAR mice after 2 weeks of rapamycin or vehicle treatment (5 weeks after HIV infection). Each dot represents an individual mouse; horizontal bars indicate median values. Mann-Whitney *U* test. \**P* < 0.05, \*\**P* < 0.01, \*\*\**P* < 0.001, \*\*\*\**P* < 0.0001.

We previously demonstrated that low-dose and intermittent rapamycin treatment decreases chronic inflammation and improves anti-viral T cell functions in HIV infected humanized mice (30). To further evaluate whether rapamycin treatment alleviates HSCs derived CAR T exhaustion and restores control over viral replication, we treated HIV infected mice with either rapamycin or vehicle for two weeks at 0.5mg/kg, 3 days a week as described in (30). Compared to vehicle-treated mice, we observed a significant decrease in the activation markers HLA-DR and CD38, and exhaustion markers PD-1 and Tim-3, among CAR T cells in the peripheral blood (**Fig. 2G**). Most importantly, we observed lower viral loads in rapamycin-treated CAR mice compared to either mock or vehicle-treated CAR mice (**Fig. 2H**). Taken together, our data suggest that rapamycin treatment alleviates CAR T exhaustion and potentially restores the anti-viral functions of CAR T cells.

### Transcriptomic changes of CAR-T cells following rapamycin treatment in HIV infected humanized mice

It is evident that rapamycin treatment changes the metabolic and functional activities of T cells *in vivo*. To more closely examine the effects of rapamycin treatment and how it can affect the CAR-T cell transcriptome, HIV infected HSCs-CAR humanized BLT mice were either treated with rapamycin or vehicle for two weeks before necropsy. Afterward, splenocytes were isolated and CAR+ cells were sorted (based on GFP expression), and bulk RNA sequencing was performed (schematically shown in **Fig.3A**). A total of 0.5-1 million CAR (GFP+) cells were sorted as shown in **Fig. 3B** from each mouse. Principal component analysis (PCA) showed that CAR cells derived from untreated mice clustered separately from CAR cells isolated from rapamycin treated mice (**Fig. 3C**). Notably, as shown in **Fig. 3D**, the heatmap and corresponding dendrogram clusters highlighted a significant downregulation of exhaustion-related markers, including inhibitory receptors PDCD1 (PD-1), Tim3, LAG3, SLAM7, and exhaustion transcription factors EOMES and TOX (33–38) in the rapamycin-treated CAR cells. Furthermore, rapamycin-induced a decrease in type I interferon-related genes, such as CXCL13 and IRFs (IRF1, IRF4), which are known contributors to T-cell exhaustion (39–42). Corroborating our cytometric analysis (**Fig. 2G**), these changes suggest that rapamycin mitigates T-cell exhaustion.

**Figure 3.**
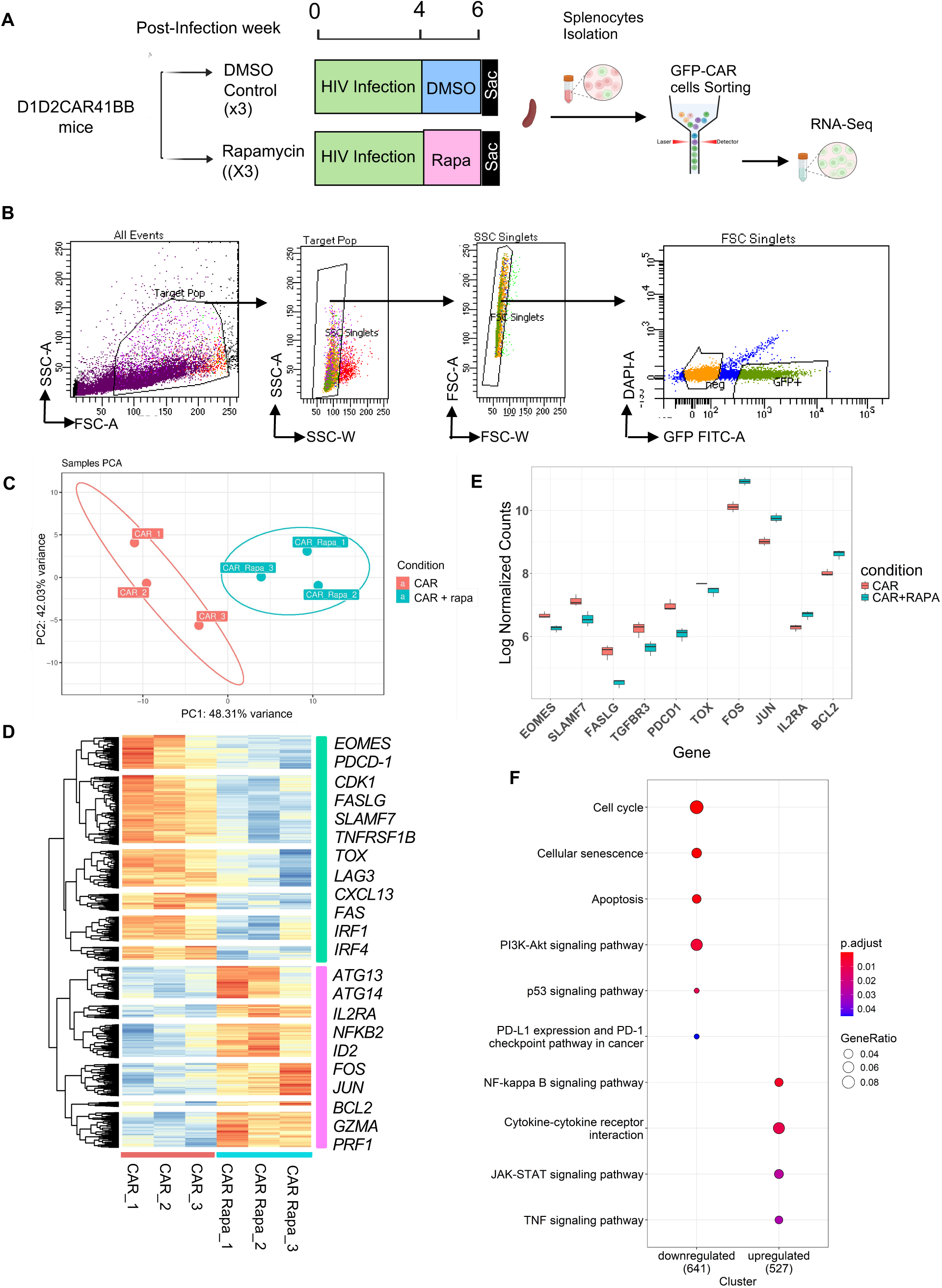
Transcriptional signatures showed reduced exhaustion markers and upregulated of memory and survival related signaling in CAR T cells in rapamycin treated mice. **A)** Humanized NSG-BLT mice with D1D2CAR41BB modified HSCs were treated with rapamycin or vehicle for 2 weeks before necropsy. Afterwards, splenocytes were isolated and GFP+ CAR cells were sorted, and bulk RNA sequencing was performed. **B)** Representative flow cytometry analysis showing the gating strategy for sorting of GFP+ CAR single cells. **C)** Principal component analysis and **D)** Heatmap showing the relative expression (z score) of the top 5,000 genes that were differentially expressed between the 2 populations of CAR T cells derived from rapamycin-treated versus vehicle-treated CAR mice. Genes were divided into downregulated (Green) and upregulated (Pink) clusters by K-means clustering based on expression. **E)** Boxplot of Log normalized counts of genes important in T cell survival, activation, and exhaustion. **F)** KEGG pathway analysis of differentially expressed genes among CAR T cells between rapamycin-treated and vehicle-treated CAR mice. GeneRatio‘Gene ratio’ is the percentage of total DEGs in the given GO term.

Intriguingly, rapamycin treatment also led to an upregulation of genes associated with T-cell survival and persistence, notably those of the activator-protein-1 (AP-1) family, such as JUN and FOS (43–45) as shown in **Fig. 3D**. As rapamycin is a potent autophagy inducer (30, 46), autophagy-related genes such as ATG13 and ATG14 were also elevated post-treatment. Moreover, genes such as IL2R, GZMA and PRF1, encoding IL2 receptor, granzyme A and perforin, respectively, which are crucial for the cytolytic activity of T cells, displayed increased expression in the rapamycin-treated CAR cells (**Fig. 3D**). The box plot visualization revealed significant differences in the log-normalized counts of key genes between the two conditions as shown in **Fig.3E**.

We further investigated the gene expression involved in biological pathway signaling using KEGG pathway analysis shown in **Fig. 3F**. Downstream targets of mTOR, including several cyclin-dependent kinases (CDKs) that are involved in cell cycling pathways, were reduced. Concurrently, cell senescence and apoptotic pathways were downregulated, as shown by lower FAS signaling and increased expression of the anti-apoptotic gene BCL2 in rapamycin-treated CAR T-cells (**Fig. 3F and Fig. 3E**). At the same time, NF-kB, JAK-STAT, and TNF signaling pathways were upregulated in CAR T cells from rapamycin treated mice. Collectively, these findings underscore rapamycin’s comprehensive role in modulating CAR T-cell function, effectively reducing exhaustion markers, and enhancing both survival and cytotoxic capabilities.

### Low-dose rapamycin treatment in combination with ART reduced exhaustion of CAR T cells and significantly reduced viral rebound

To further assess the effects of rapamycin treatment in combination with ART, we treated HIV infected humanized mice transplanted with mock or CAR-modified HSCs with rapamycin or vehicle for 2 weeks, followed by 4 weeks of ART treatment and ART withdrawal (**Fig 4A**). As shown in **Fig. 4B**, compared to control mice, ART combined with rapamycin significantly reduced the expression of immune activation markers (HLA-DR, CD38), and exhaustion markers (PD-1, Tim3) among CAR-T cells in the blood and spleen at necropsy. TOX is a key transcription factor that regulates the T cell exhaustion program and the expression of TOX has been associated with cellular exhaustion during chronic infection (33, 35, 47). Similar to our RNAseq studies (**Fig 3D**), flow analysis of CAR-T cells at necropsy also showed significant down regulation of TOX expression of CAR-T cells from mice treated with rapamycin as compared to vehicle control (**Fig. 4C**).

**Figure 4:**
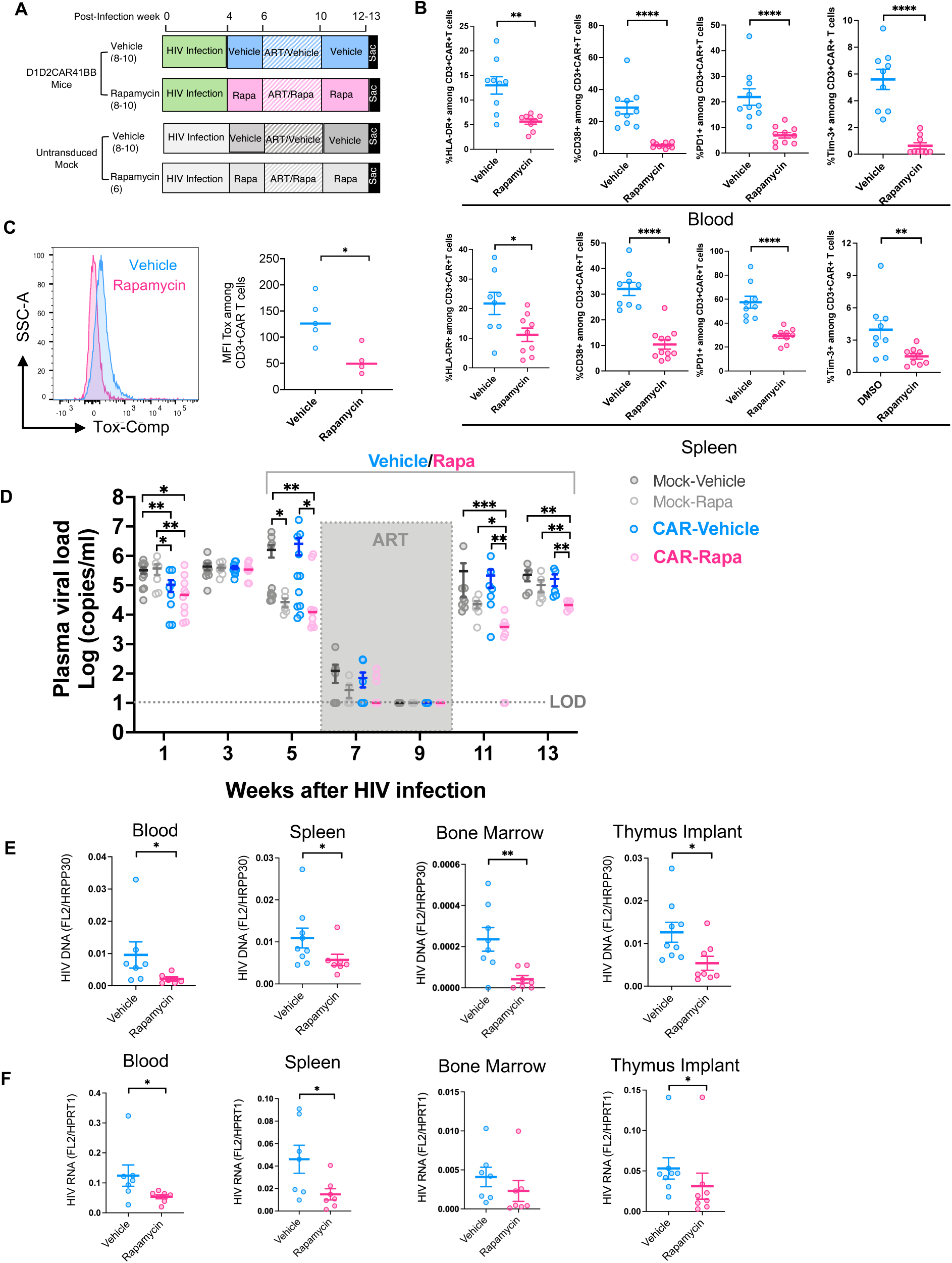
Long-term low dose rapamycin treatment in combination with ART alleviated CAR-T cell exhaustion and reduced viral rebound. **A)** Humanized NSG-BLT mice with either D1D2CAR41BB modified or non-modified HSCs were infected with HIV and treated with rapamycin or vehicle for 2 weeks. Afterwards, while continuing rapamycin or vehicle treatment, mice were treated with ART for 4 weeks, followed by ART interruption for 2-3 weeks. (**B**) PD-1, Tim-3, HLA-DR and CD38 expression was measured by flow cytometry (quantified by gating percentage positive cells) on peripheral blood(top) or spleen(bottom) CD3 CAR+T cells. **C)** Splenocytes from CAR mice treated with ART and rapamycin or vehicle were isolated and stained with intracellular Abs against human Tox1. MFIs of the Tox1 on CAR+ T cells were measured by flow cytometry. **D)** Longitudinal HIV viral load in plasma from humanized mice after rapamycin or vehicle treatment were measured by real-time PCR. Dotted line indicates limit of detection. **E)** HIV DNA copies per cell from PBMC, splenocytes, bone marrow or thymus implant from CAR transduced mice as measured by real-time PCR. Human HRPP30 gene was used as internal control. **F)** Relative HIV cellular RNA expression from multiple lymphoid tissues from CAR transduced mice after the indicated treatment. Human HPRT1 gene expression was used as internal control (*n* = 5–7 per group). Mann-Whitney test (unpaired); \**P* < 0.05, \*\**P* < 0.01, \*\*\**P* < 0.001, \*\*\*\**P* < 0.0001.

To study the effects of rapamycin treatment on HIV suppression, persistence, and viral rebound in CAR mice, plasma viral loads were measured longitudinally. As shown in **Fig. 4D**, at 9 weeks post-HIV infection and after 4 weeks of ART, all mice exhibited full viral suppression with undetectable viral load. One week after interrupting ART, all mock control and vehicle-treated CAR mice, and all but 2 rapamycin-treated CAR mice experienced viral rebound. Three weeks after ART withdrawal, all mice showed viral rebound. However, rapamycin-treated CAR mice maintained a significantly lower viral load as compared to vehicle-treated CAR mice, or mock mice treated with either vehicle or rapamycin. Additionally, we observed significantly lower levels of viral DNA (**Fig. 4E**) and cell associated HIV RNA (**Fig. 4F**) in the blood, spleen, and bone marrow at necropsy after ART withdrawal, indicating a reduction in overall viral burden in the rapamycin-treated CAR mice. Taken together, these data suggest that the combination of rapamycin and ART treatment improves CAR T cell function-in HIV suppression and reduces viral rebound.

### Rapamycin treatment reduced mitochondria ROS in CAR T cells and improved CAR T function

Excessive ROS can cause damage to cellular components, including lipids, proteins, and DNA (48, 49). Decreased mitochondrial biogenesis and excessive production of mitochondrial ROS can exacerbate mitochondrial dysfunction and immune exhaustion in T cells (50–54). Our *in vitro* data suggested that rapamycin treatment protects the mitochondria against oxidative stress in CAR T cells (**Fig 1D**). To further investigate the potential for *in vivo* rapamycin treatment to reduce oxidative stress and alleviate mitochondrial injuries in CAR T cells, we measured mitochondrial dysfunction levels using MitoSOX to detect mitochondrial superoxide levels in CAR T cells from both peripheral blood and spleen of CAR mice treated with rapamycin or vehicle control. We observed that CAR T cells from mice treated with rapamycin exhibited significantly lower mitochondrial ROS compared to vehicle-treated mice (**Fig. 5A**), suggesting a reduction of oxidative stress in treated mice.

**Figure 5.**
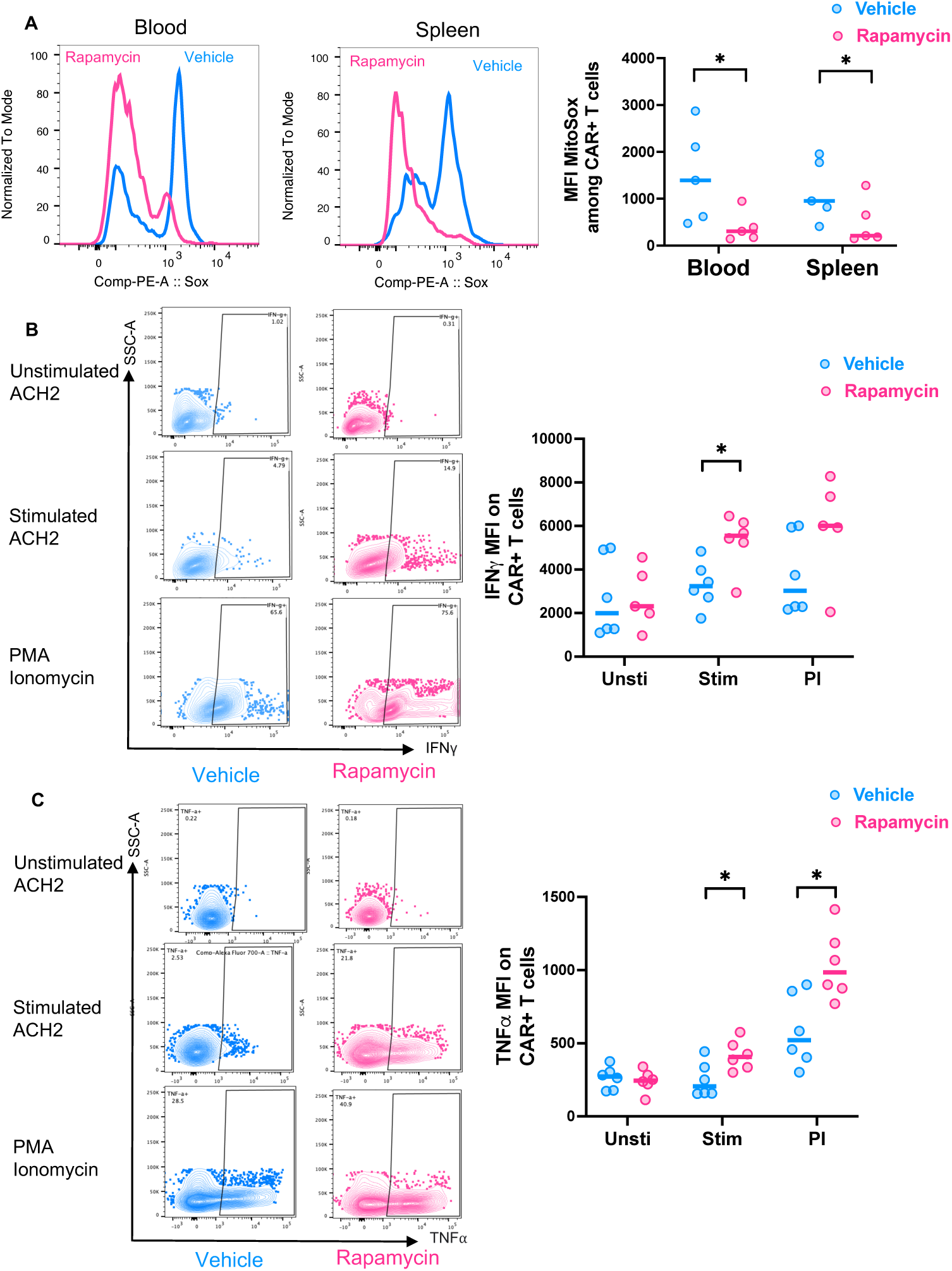
Rapamycin treatment decreases mitochondria ROS and improves CAR T function *in vivo*. **A)** ROS levels analyzed by MitoSOX staining in blood or spleen CAR T cells isolated from CAR mice treated with either rapamycin or vehicle. **B-C)** Splenocytes from CAR transduced, HIV-1–infected, vehicle-treated, or rapamycin-treated mice were stimulated with PMA/ionomycin, or envelope expressing (stimulated ACH2) or non-expressing (unstimulated ACH2) cells and production of IFN-γ and TNF-⍺ by CAR T cells was measured by flow cytometry. Representative flow cytometry data showing percentage and MFI of IFN-γ^+^ **(B)** and TNF-⍺ **(C)** among CAR T cells from HIV-1–infected, vehicle-treated, or rapamycin-treated mice. Each dot represents an individual mouse; horizontal bars indicate median values. Mann-Whitney test (unpaired); \**P* < 0.05, \*\**P* < 0.01.

To further investigate whether HIV-specific CAR T cell responses were improved in the ART and rapamycin combined-treatment group, splenocytes from rapamycin or vehicle treated mice were stimulated with mitogens PMA/ionomycin, or stimulated with HIV target cells (stimulated latently infected ACH2 cells, which are Env+) or control cells (unstimulated ACH2 cells, which are Env-). Compared with the vehicle-treated control, CAR T cells from rapamycin-treated infected mice produced significantly higher levels of pro-inflammatory IFN-γ and TNF-α cytokines after PMA/ionomycin stimulation (**Figs. 5B, 5C**), and HIV Env+ cells, indicating improved proinflammatory cytokine production and anti-viral responses of CAR T cells. In summary, these data suggest that a combination of rapamycin and ART reduced ROS and improved anti-HIV functions of CAR T cells *in vivo*.

### Short-term Rapamycin treatment in CAR-HSCs NSG-Tg (IL-15) humanized mice showed delayed viral rebound and smaller reservoirs after ART withdraw

An enhanced humanized mouse model, termed Hu-NSG-Tg(IL-15), which was engineered to express physiological level of human IL-15, was reported to support more robust engraftment of human immune cells, including T cells, B cells, and NK cells, and therefore represents a valuable model for studying HIV pathogenesis and immune responses (55, 56). We adopted this new model to make HSCs-CAR mice and examined the effects of rapamycin or vehicle treatment. Importantly, to examine if the effects of rapamycin treatment persist after short-term treatment, rapamycin treatment was started with ART and stopped 2 weeks after ART withdrawal, and mice were continuously monitored for an additional 3 weeks before necropsy (shown in **Fig 6A**). As shown in **Fig. 6B**, two weeks after ART interruption, viral loads quickly rebounded in all mice (100%, 4/4) from the mock group. In CAR-HSCs vehicle-treated group, two out of four did not rebound (50%, 2/4) 2 weeks after ART withdrawal. However, all mice rebounded 4 weeks after ART interruption. In contrast, none of the rapamycin-treated CAR-HSCs mice had viral rebound 2 weeks after ART. 2 weeks after cessation of rapamycin treatment and 4 weeks after ART cessation, three out of six (50%) rapamycin CAR-HSCs mice did not rebound. 5 weeks after ART withdrawal at necropsy, two out of six (33%) rapamycin-treated CAR-HSCs mice continued to have undetectable viral load, while the other four maintained significantly lower viral loads after rebound. In summary, we observed that rapamycin-treated CAR mice had significantly improved CAR suppression of viral replication, leading to significantly delayed viral rebound after ART cessation, as shown in **Fig 6C**. Importantly, As shown in **Fig 6D**, we observed an overall decrease in the level of cell associated HIV DNA and RNA in the spleen and bone marrow in rapamycin-treated CAR mice compared to vehicle-treated CAR mice, and mice with undetectable viral loads also showed low/undetectable viral DNA/RNA in tissues, suggesting lower reservoir in rapamycin-treated CAR mice.

**Figure 6.**
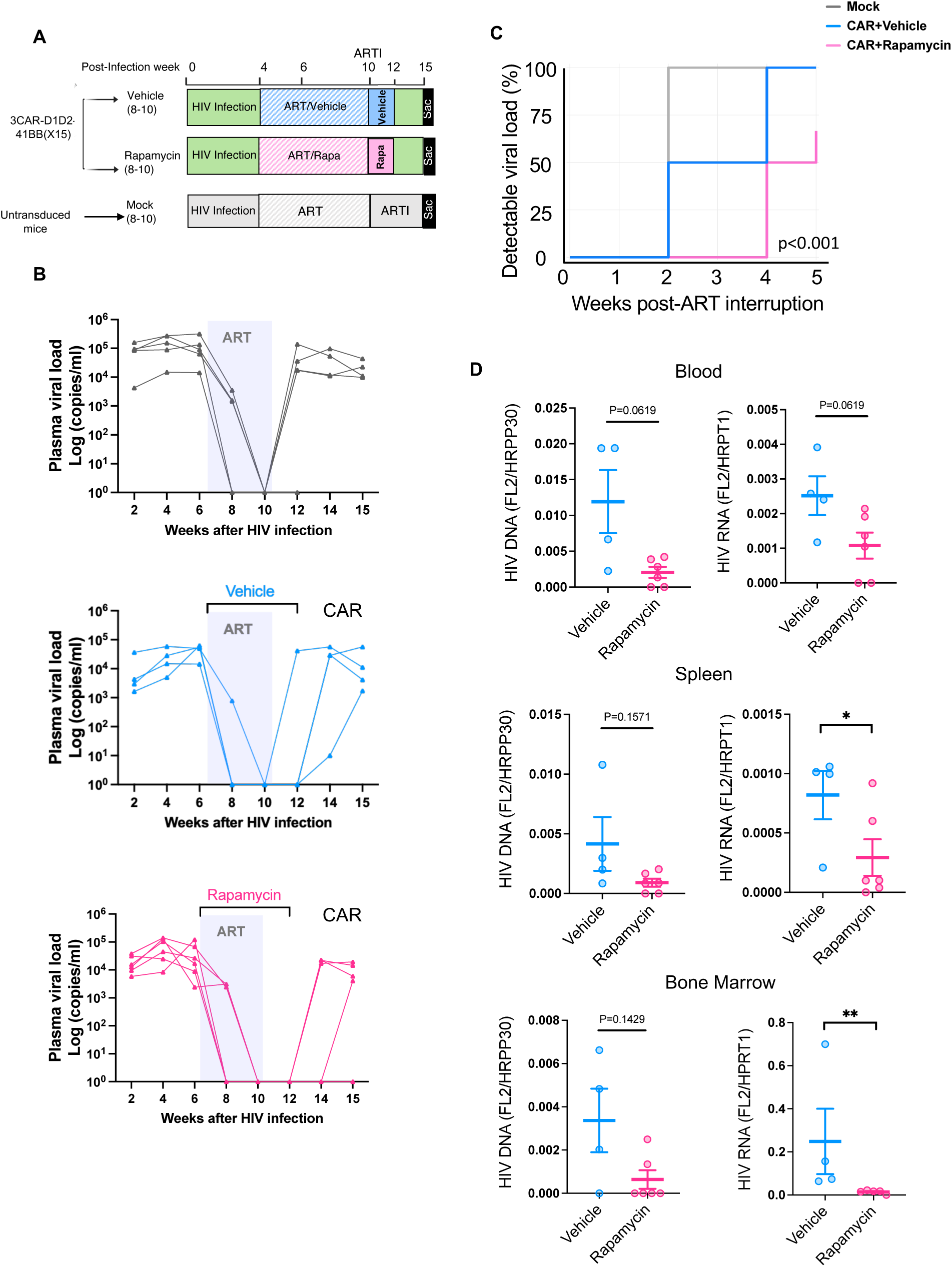
Rapamycin treated NSG-IL-15 CAR mice showed significantly delayed viral rebound and smaller reservoirs after ART withdraw. **A)** Humanized NSG-IL15-BLT mice were constructed with either unmodified HSCs or HSCs modified with D1D2CAR41BB. After immune reconstitution, mice were infected with HIV-1_NFNSXL9_. Four weeks after infection, mock mice were treated with ART only. Mice with CAR modified HSCs were treated with rapamycin or vehicle along with ART. Following successful viral load suppression, ART was interrupted and rapamycin or vehicle treatment continued for two additional week before discontinuation. **B)** Longitudinal plasma HIV viral load as measured by real-time PCR. Dotted line indicates limit of detection. **C)** Survival analysis of time to detectable viral load among mock, CAR mice that were treated with vehicle or rapamycin. p<0.0001 by log-rank test. **D)** HIV DNA and relative cellular HIV RNA expression from blood PBMCs, splenocytes and bone marrow as measured by real-time PCR. Mann-Whitney test (unpaired); *P < 0.05, **P < 0.01.

## DISCUSSION

CAR redirected T cell immunity against HIV-1 represents a highly promising approach that can be used in most HIV infected individuals. CARs recognize target cells through direct binding to specific cell surface antigens and are HLA-unrestricted, bypassing a major limitation for T cell immunotherapies (57). Now widely applied for cancer, some of the first CAR clinical trials were for HIV-1 infection (58–60). The “original” HIV-specific CARs were composed of a CD4 extracellular domain linked to the intracellular CD3-ζsignaling domain (CD4CAR), utilizing CD4 binding to HIV-1 Envelope (Env) for targeting and killing of HIV-1-infected cells (8). HIV-1 envelope interaction with CD4 is critical for viral replication, thus limiting viral escape from a CD4-based CAR (61). Tremendous progress has been made using CD4 based CAR T cell therapy against HIV infection since its invention. Improvement in the CAR design have been made with modification of the Env recognition domain and inclusion of co-stimulatory domains, and enhanced CD4-based CAR-T cell efficacy has been demonstrated by multiple groups of investigators (6, 15, 62–64), including ours (10, 12, 13, 65–68), and it is currently under investigation in multiple clinical trials (ClinicalTrials.gov Identifier: NCT03617198, NCT04648046).

Given the challenge of HIV latent reservoirs driving chronic infection and persistence under suppressive ART, the functional persistence of CAR T cells is critical for successful long-term immune containment of HIV infection. HSCs based gene therapy supports lifelong generation of functional immune progeny, giving rise to a stable supply of gene-modified CAR T cells. We have shown that HSCs-based CAR therapy allows for lifelong, persistent production of functional CAR-T cells to control viral replication from reactivated reservoirs, and HSCs-based CAR therapy showed better persistence and trafficking to tissue reservoirs than periphery blood derived CAR T cells (10, 11, 13, 65, 67). Despite their *in vivo* efficacy in reducing viral replication and reservoir, HSCs-derived CAR-T cells cannot achieve full viral suppression after ART withdrawal. Persistent type I Interferon signaling and immune activation during chronic HIV infection are driving forces of immune dysfunction (69, 70), and engineered CAR-T cells are also subject to this immune exhaustion. Previously we demonstrated that targeting persistent type I interferon signaling, either by blocking type I interferon receptor (71), or inducing autophagy by rapamycin (30, 69, 72), can lead to reduced hyperinflammation and improved anti-HIV T cell function *in vivo*. In the current study, we found that rapamycin treatment also significantly improved CAR-HSCs therapy in HIV infected humanized mice. Rapamycin treatment reduced persistent immune activation, rejuvenated CAR-T cell function, leading to delayed viral rebound and better viral control after ART cessation, even after rapamycin treatment was stopped. Notably, two out of six rapamycin treated CAR-HSCs continued to have undetectable viral loads five weeks after ART withdrawal and three weeks after rapamycin cessation, while all other groups had 100% viral rebound. The rapamycin treatment also led to reduced cell-associated HIV RNA and DNA in blood and multiple lymphoid tissues in CAR-HSCs mice. Importantly, we observed improved mitochondria function and significant transcriptomic modification of CAR-T cells by rapamycin treatment, suggesting the dual beneficial effects of rapamycin in reducing T cell inhibitory receptors while simultaneously promoting stemness-related gene expression, offering promising insights into the optimization of CAR T-cell therapy for sustained anti-HIV responses.

Rapamycin is a mTOR inhibitor, which is a major regulator of cellular metabolism and the cellular ageing process. First discovered and FDA-approved for treatment of various cancers and as an immunosuppressant, rapamycin’s effects on aging is increasingly recognized and used for longevity studies (20, 73). Multiple studies suggest that rapamycin treatment extends health span and improves the function of the aging immune system, such as improving antiviral activities in older adults (21, 22, 28). This is of relevance to HIV, as people living with HIV are aging with greater life expectancy due to effective ART. Despite the success of ART, the difference in comorbidity-free years between PLWH and the general population persists, and premature immune aging has been shown to be the major culprit driving age-related co-morbidity (74–76). Treatment with Rapamycin was correlated with reduced HIV reservoir in HIV-1 infected kidney transplant recipients (77). However, as a master regulator, mTOR signaling plays a key role in T cell fate, such as effector versus memory differentiation, and its regulation needs to be tightly controlled (78). Treatment with daily rapamycin can limit T cell proliferation in SIV infected rhesus macaques on antiretroviral therapy and was not shown to impact SIV reservoir (79). For our current study, we chose to use lower and intermittent dosing of rapamycin that we previously described, which did not impact T cell homeostasis (30). Therefore, careful studies on the dosing and treatment regimen of rapamycin are needed to maximize its effects and reducing its toxicity in animal models and in clinical studies.

In summary, our study describes the effects of rapamycin on improving the function of anti-HIV CAR-T cells *in vivo*, demonstrating its impact on CAR-T cell metabolism and transcriptomic modification. We believe that the results described in this study shed light on potential strategies to augment CAR-T functions for treating HIV infection, and our findings may also be applicable to other CAR-T therapies that are affected by immune exhaustions (80).

## MATERIALS AND METHODS

### Lentivirus production

The lentivirus-based D1D2CAR41BB vectors were produced in Lenti-X 293T cells using the Lipofectamine 2000 reagent (Invitrogen). Briefly, Lenti-X 293T cells were co-transfected with D1D2CAR41BB vector with pCMV.ΔR8.2.Δvpr packaging construct and the pCMV-VSV-G envelope protein plasmid, as previously described (1, 2). The supernatant was obtained from transfected Lenti-X 293T cells 48 hours post-transfection. It underwent filtration using a 0.45 μm sterile filter and concentration through ultracentrifugation using a Beckman SW32 rotor at 154,000g at 4°C. Following aspiration of the medium, the pellet was resuspended in PBS and stored at –80°C.

### Transduction of CAR T cells

Primary T cells were sorted from primary PBMC from healthy donors using Pan T cell isolation kit – human (Miltenyi Biotec, # 130-096-535). Isolated T cells were stimulated with plate-bound anti-CD3 and anti-CD28 (Miltenyi Biotec) at 2 million cells/ml. After activation for at least 24 hrs, cells were washed and transduced with CD4CAR vector on retronectin-coated plate with cytokine IL-2. Cells were cultured in RPMI supplemented with 10% FBS and 1% Pen/strip with IL-2 for additional 2 weeks. Following this culture period, the cells were treated with either DMSO control or rapamycin at a concentration of 500pM for 2 days, prior to conducting the Seahorse assay.

### Seahorse assay

250,000 CAR-T cells were seeded into wells of a poly-d-lysine-coated (100 μg/mL) XF96 spheroid plate. Cells in the plate were centrifuged at 450 rpm for 7 min with no centrifuge brake, mitochondrial respiration was measured using the Seahorse XF96 extracellular flux analyzer equipped with a spheroid plate-compatible thermal tray (Agilent Technologies). Basal respiration was first measured in 3 mM glucose media. To validate cell respirometry with the XF96 spheroid plate, CAR-T cells were then sequentially exposed to glucose (final concentration in well of 20 mM), Oligomycin A (3.5–4.5 μM final concentration), FCCP (1 μM final concentration) and Antimycin A (Ant A, 2.5 μM final concentration).

### Humanized mice generation

D1D2CAR41BB BLT mice were constructed similarly to previously reported HIV-1 Triple CAR BLT humanized mice(68). Briefly, human fetal liver derived CD34+ cells were purified by immunomagnetic separation. Cells were then transduced overnight with D1D2CAR41BB lentiviruses with retronectin-coated plates. On day of transplant, NOD.Cg-PrkdcscidIl2rgtm1Wjl/SZJ (NOD/SCID/IL2Rγ−/− or NSG, The Jackson Laboratory) or NOD.Cg-Prkdcscid Il2rgtm1Wjl Tg(IL15)1Sz/SzJ (NSG-huIL15, The Jackson Laboratory) mice received 2.7 Gy total body sublethal irradiation and then were transplanted with transduced CD34+ in Matrigel (Corning Life Sciences), liver and thymus tissue under the kidney capsule, with tissue from the same donor as the CD34+ cells. Afterward, mice were injected with ∼0.5×10^6^ lentivirus-based CAR vector transduced CD34+ cells. At 8–10 weeks post-transplantation, each mouse was bled retro-orbitally and peripheral blood mononuclear cells analyzed by flow cytometry to check human immune cell engraftment. Upon stable human leukocyte reconstitution efficiency more than 50%, mice were used for HIV-1 infection and further experiments.

### HIV-1 infection, ART and rapamycin treatment

The R5 tropic strain of HIV-1_NFNSXSL9_ was generated by transfection of 293T cells with plasmid containing full-length HIV-1_NFNSXSL9_ genome. Humanized mice were infected with HIV-1_NFNSXSL9_ (500 ng p24 per mouse) through retro-orbital injection while under inhalant general anesthesia. Infected mice with demonstrable viral infection were treated for 6 weeks with ART drugs. The ART regimen is consisted of tenofovir disoproxil-fumarate (TDF, 80mg/kg), emtricitabine (FTC, 120mg/kg), and Elvitegravir (ELV160mg/kg) given by food. TDF, FTC and ELV were generously supplied by Gilead Sciences. TDF, FTC and ELV were dissolved in DMSO and mixed with sweetened moist gel meal (DietGel Boost, ClearH2O; Medidrop Sucralose) as previously described (81). For rapamycin treatment, mice were injected i.p. with 0.5 mg/kg rapamycin (LC laboratories) 3 times a week.

### Flow cytometry

Mitochondria-associated ROS levels were measured by staining cells with MitoSOX (Molecular Probes/Invitrogen) at 5 μM for 40 min at 37 °C. Cells were then washed with PBS solution and resuspended in PBS solution containing 2% FBS for FACS analysis. Single-cell suspensions prepared from peripheral blood or spleen of humanized mice were stained using the following antibodies for flow cytometry: CD45 (clone HI30), CD3 (clone OKT3), CD4 (clone RPA-T4), CD8 (clone SK1), CD38 (clone HIT2), HLA-DR (clone L240), CD45RA (clone HI100), CD62L (clone DREG-56), IFN-γ (clone 4S.B3), IL-2 (clone MQ1-17H12), TNF-α (clone Mab11), Tox (clone TXRX10), PD-1 (clone ebioJ105), and, TIM-3 (clone F38-2E2). LIVE/ DEAD Fixable Yellow Dead Cell Stain Kit (Invitrogen) was used. Abs for cell surface markers and intracellular markers were conjugated to FITC, phycoerythrin (PE), PerCP-Cy5.5, PE-Cy5, PE-Cy7, electron coupled dye (ECD), allophycocyanin (APC), APC–eFluor780, Alexa Fluor 700, eFluor450, Pacific Orange, or Pacific Blue in the appropriate combination. The LSRFortessa flow cytometer and FACSDiva software (BD Biosciences) were employed to obtain the cells, while FlowJo software was used for data analysis. A minimum of 1000 cells were acquired for each analysis, and each flow plot, representative of the data, was replicated more than three times.

### Nucleic acid extraction and real time PCR

To measure HIV plasma viremia, viral RNA was extracted from plasma and 1-step real-time PCR was performed using the TaqMan RNA-to-Ct 1-Step Kit (Thermo Fisher Scientific, USA) with the following primers and probe:

HIV-1 forward primer: 5′-CAATGGCAGCAATTTCACCA-3′;
HIV-1 reverse primer: 5′-GAATGCCAAATTCCTGCTTGA-3′;
HIV-1 probe: 5′-[6-FAM] CCCACCAACAGGCGGCCTTAACTG [Tamra-Q]-3′;

To measure the levels of cell-associated HIV RNA with HPRT1 as an internal control, cells were harvested for RNA extraction according to manufacturer’s protocol (Qiagen) and making of cDNA using the High-Capacity cDNA Reverse Transcription Kit (Thermo Fisher Scientific). For HPRT1, Single Tube TaqMan Gene Expression Assays (Thermo Fisher Scientific) human HPRT1 (Hs01003267_m1) was used. Relative mRNA expression was calculated by normalizing genes to HPRT1 mRNA expression.

To measure the levels of cell-integrated HIV DNA with RPP30 as an internal control, cells were harvested for DNA extraction according to manufacturer’s protocol (Qiagen). For HIV DNA as well as RPP30, Single Tube TaqMan Gene Expression Assays (Thermo Fisher Scientific). Relative DNA integration was calculated by normalizing HIV DNA to RPP30 gene count.

HIV-1 forward primer: 5′-CAATGGCAGCAATTTCACCA-3′;
HIV-1 reverse primer: 5′-GAATGCCAAATTCCTGCTTGA-3′;
HIV-1 probe: 5′-[6-FAM] CCCACCAACAGGCGGCCTTAACTG [Tamra-Q]-3′;
RPP30 Forward primer: 5′-GATTTGGACCTGCGAGCG-3′;
RPP30 Reverse primer: 5′-GCGGCTGTCTCCACAAGT-3′;
RPP30 Probe: /5HEX/TTCTGACCT/ZEN/GAAGGCTCTGCG/3IABkFQ/-3′

### D1D2CAR sorting and RNA sequencing

Spleens from D1D2CAR transduced mice were collected, mashed over a 70 μm cell strainer, and resuspended into single cell suspensions in complete RPMI after red blood cell lysis using ACK lysis buffer (Thermo Fisher). 0.5-1 million GFP+ single live cells were sorted using BD FACS Melody (BD Biosciences) from splenocytes per mouse. Three replicates of each experiment were carried out. RNA extraction was done using RNeasy kits (Qiagen). Sample QC and integrity (RIN-equivalent values) was performed using Tapestation Analysis software v3.2, Agilent Technologies. Sequencing was carried out using Illumina NovaSeq platform. Raw sequence data of different treatment conditions (in triplicate) were pre-processed for quality using Fastqc. Trimmomatic was used for adaptors and quality trimming. After this, reads were aligned onto human genome (hg38) using STAR aligner. SAMtools was used to convert SAM files BAM files. Mapped reads were counted across human genes by using tool featureCounts (75) that provided raw counts data by assigning mapped reads to genes. Differential gene expression analysis with the raw read counts data using R package DESeq2. Raw sequence data and processed data have been deposited to GEO. Gene expression data was analyzed using Gene Set Enrichment Analysis (GSEA) software tool. For pathway analysis, STRING (Search Tool for the Retrieval of Interacting Genes/Proteins) software was used.

## Acknowledgement

We thank Romas Geleziunas and Jeff Murry and the people at Gilead for providing the antiretroviral drugs used in this study. We thank UCLA Center for AIDS Research (CFAR) Humanized Mouse Core staff research associate Nianxin Zhong for his assistance in the humanized mice work. This work was funded by the National Institute of Allergy and Infectious Diseases (grants 1R21AI140866 to AZ, R01AI172727 to AZ and MDM), the National Institute on Drug Abuse (grant R01DA-52841 to AZ), the American Foundation for AIDS Research (grant 110304-71-RKRL, 10395-72-RPRL to AZ and 109577-62-RGRL and 110038-67-RSRL to SGK), the National Cancer Institute (grant 1R01CA239261-01 to SGK), National Institute of Health grants P30AI28697 (to the UCLA CFAR Virology Core, Gene and Cell Therapy Core, and Humanized Mouse Core) and U19AI149504 (to SGK and Dr. Irvin Chen [UCLA]); the California Institute for Regenerative Medicine (grant TRAN1-14625 to SGK.); California HIV/AIDS Research Program (grant H24BD7817 to WM). This work was also supported by the UCLA AIDS Institute, the James B. Pendleton Charitable Trust, and the McCarthy Family Foundation. The graphic figure was created with BioRender.com.

## Author contribution

WM and AZ designed the experiments. WM, ST, JH, NK, VR, EC, VP, MAC, HM, HG and LW conducted the experiments. WM, JH, ST, NK and AZ analyzed the data. WM and AZ wrote the original draft, JH, ST, NK, HM, MDM and SGK reviewed and edited the manuscript.

## References

1. Collins DR, Gaiha GD, and Walker BD. CD8(+) T cells in HIV control, cure and prevention. Nat Rev Immunol. 2020;20(8):471–82.

2. Perdomo-Celis F, Taborda NA, and Rugeles MT. CD8(+) T-Cell Response to HIV Infection in the Era of Antiretroviral Therapy. Front Immunol. 2019;10:1896.

3. Deng K, Pertea M, Rongvaux A, Wang L, Durand CM, Ghiaur G, et al. Broad CTL response is required to clear latent HIV-1 due to dominance of escape mutations. Nature. 2015;517(7534):381–5.

4. Blackburn SD, Shin H, Haining WN, Zou T, Workman CJ, Polley A, et al. Coregulation of CD8+ T cell exhaustion by multiple inhibitory receptors during chronic viral infection. Nat Immunol. 2009;10(1):29–37.

5. Marsden MD, and Zack JA. Double trouble: HIV latency and CTL escape. Cell Host Microbe. 2015;17(2):141–2.

6. Mu W, Carrillo MA, and Kitchen SG. Engineering CAR T Cells to Target the HIV Reservoir. Front Cell Infect Microbiol. 2020;10:410.

7. Louie RHY, Kaczorowski KJ, Barton JP, Chakraborty AK, and McKay MR. Fitness landscape of the human immunodeficiency virus envelope protein that is targeted by antibodies. Proc Natl Acad Sci U S A. 2018;115(4):E564–E73.

8. Yang OO, Tran AC, Kalams SA, Johnson RP, Roberts MR, and Walker BD. Lysis of HIV-1-infected cells and inhibition of viral replication by universal receptor T cells. Proceedings of the National Academy of Sciences of the United States of America. 1997;94(21):11478–83.

9. Rafiq S, Hackett CS, and Brentjens RJ. Engineering strategies to overcome the current roadblocks in CAR T cell therapy. Nat Rev Clin Oncol. 2020;17(3):147–67.

10. Zhen A, Carrillo MA, Mu W, Rezek V, Martin H, Hamid P, et al. Robust CAR-T memory formation and function via hematopoietic stem cell delivery. PLoS Pathog. 2021;17(4):e1009404.

11. Zhen A, Kamata M, Rezek V, Rick J, Levin B, Kasparian S, et al. HIV-specific Immunity Derived From Chimeric Antigen Receptor-engineered Stem Cells. Mol Ther. 2015;23(8):1358–67.

12. Zhen A, Peterson CW, Carrillo MA, Reddy SS, Youn CS, Lam BB, et al. Long-term persistence and function of hematopoietic stem cell-derived chimeric antigen receptor T cells in a nonhuman primate model of HIV/AIDS. PLoS Pathog. 2017;13(12):e1006753.

13. Carrillo MA, Zhen A, Mu W, Rezek V, Martin H, Peterson CW, et al. Stem cell-derived CAR T cells show greater persistence, trafficking, and viral control compared to ex vivo transduced CAR T cells. Molecular therapy : the journal of the American Society of Gene Therapy. 2024;32(4):1000–15.

14. Pauken KE, and Wherry EJ. Overcoming T cell exhaustion in infection and cancer. Trends Immunol. 2015;36(4):265–76.

15. Maldini CR, Claiborne DT, Okawa K, Chen T, Dopkin DL, Shan X, et al. Dual CD4-based CAR T cells with distinct costimulatory domains mitigate HIV pathogenesis in vivo. Nature medicine. 2020;26(11):1776–87.

16. Rust BJ, Kean LS, Colonna L, Brandenstein KE, Poole NH, Obenza W, et al. Robust expansion of HIV CAR T cells following antigen boosting in ART-suppressed nonhuman primates. Blood. 2020;136(15):1722–34.

17. Grosser R, Cherkassky L, Chintala N, and Adusumilli PS. Combination Immunotherapy with CAR T Cells and Checkpoint Blockade for the Treatment of Solid Tumors. Cancer Cell. 2019;36(5):471–82.

18. Martins F, Sofiya L, Sykiotis GP, Lamine F, Maillard M, Fraga M, et al. Adverse effects of immune-checkpoint inhibitors: epidemiology, management and surveillance. Nat Rev Clin Oncol. 2019;16(9):563–80.

19. Bajwa R, Cheema A, Khan T, Amirpour A, Paul A, Chaughtai S, et al. Adverse Effects of Immune Checkpoint Inhibitors (Programmed Death-1 Inhibitors and Cytotoxic T-Lymphocyte-Associated Protein-4 Inhibitors): Results of a Retrospective Study. J Clin Med Res. 2019;11(4):225–36.

20. Mannick JB, and Lamming DW. Targeting the biology of aging with mTOR inhibitors. Nat Aging. 2023;3(6):642–60.

21. Mannick JB, Del Giudice G, Lattanzi M, Valiante NM, Praestgaard J, Huang B, et al. mTOR inhibition improves immune function in the elderly. Science translational medicine. 2014;6(268):268ra179.

22. Mannick JB, Teo G, Bernardo P, Quinn D, Russell K, Klickstein L, et al. Targeting the biology of ageing with mTOR inhibitors to improve immune function in older adults: phase 2b and phase 3 randomised trials. Lancet Healthy Longev. 2021;2(5):e250–e62.

23. Di Benedetto F, Di Sandro S, De Ruvo N, Montalti R, Ballarin R, Guerrini GP, et al. First report on a series of HIV patients undergoing rapamycin monotherapy after liver transplantation. Transplantation. 2010;89(6):733–8.

24. Henrich TJ, Schreiner C, Cameron C, Hogan LE, Richardson B, Rutishauser RL, et al. Everolimus, an mTORC1/2 inhibitor, in ART-suppressed individuals who received solid organ transplantation: A prospective study. Am J Transplant. 2021;21(5):1765–79.

25. Wirth M, Schwarz C, Benson G, Horn N, Buchert R, Lange C, et al. Effects of spermidine supplementation on cognition and biomarkers in older adults with subjective cognitive decline (SmartAge)-study protocol for a randomized controlled trial. Alzheimers Res Ther. 2019;11(1):36.

26. Pollizzi KN, Patel CH, Sun IH, Oh MH, Waickman AT, Wen J, et al. mTORC1 and mTORC2 selectively regulate CD8(+) T cell differentiation. The Journal of clinical investigation. 2015;125(5):2090–108.

27. Pearce EL, Walsh MC, Cejas PJ, Harms GM, Shen H, Wang LS, et al. Enhancing CD8 T-cell memory by modulating fatty acid metabolism. Nature. 2009;460(7251):103-7.

28. Mannick JB, Morris M, Hockey HP, Roma G, Beibel M, Kulmatycki K, et al. TORC1 inhibition enhances immune function and reduces infections in the elderly. Science translational medicine. 2018;10(449).

29. Ando S, Perkins CM, Sajiki Y, Chastain C, Valanparambil RM, Wieland A, et al. mTOR regulates T cell exhaustion and PD-1-targeted immunotherapy response during chronic viral infection. The Journal of clinical investigation. 2023;133(2).

30. Mu W, Rezek V, Martin H, Carrillo MA, Tomer S, Hamid P, et al. Autophagy inducer rapamycin treatment reduces IFN-I-mediated Inflammation and improves anti-HIV-1 T cell response in vivo. JCI insight. 2022;7(22).

31. McLane LM, Abdel-Hakeem MS, and Wherry EJ. CD8 T Cell Exhaustion During Chronic Viral Infection and Cancer. Annual review of immunology. 2019;37:457–95.

32. Hashimoto M, Kamphorst AO, Im SJ, Kissick HT, Pillai RN, Ramalingam SS, et al. CD8 T Cell Exhaustion in Chronic Infection and Cancer: Opportunities for Interventions. Annual review of medicine. 2018;69:301–18.

33. Alfei F, Kanev K, Hofmann M, Wu M, Ghoneim HE, Roelli P, et al. TOX reinforces the phenotype and longevity of exhausted T cells in chronic viral infection. Nature. 2019;571(7764):265–9.

34. Scott AC, Dundar F, Zumbo P, Chandran SS, Klebanoff CA, Shakiba M, et al. TOX is a critical regulator of tumour-specific T cell differentiation. Nature. 2019;571(7764):270–4.

35. Khan O, Giles JR, McDonald S, Manne S, Ngiow SF, Patel KP, et al. TOX transcriptionally and epigenetically programs CD8(+) T cell exhaustion. Nature. 2019;571(7764):211–8.

36. Buggert M, Tauriainen J, Yamamoto T, Frederiksen J, Ivarsson MA, Michaelsson J, et al. T-bet and Eomes are differentially linked to the exhausted phenotype of CD8+ T cells in HIV infection. PLoS pathogens. 2014;10(7):e1004251.

37. O’Connell P, Hyslop S, Blake MK, Godbehere S, Amalfitano A, and Aldhamen YA. SLAMF7 Signaling Reprograms T Cells toward Exhaustion in the Tumor Microenvironment. Journal of immunology (Baltimore, Md : 1950). 2021;206(1):193–205.

38. Seo W, Jerin C, and Nishikawa H. Transcriptional regulatory network for the establishment of CD8(+) T cell exhaustion. Exp Mol Med. 2021;53(2):202–9.

39. Tietscher S, Wagner J, Anzeneder T, Langwieder C, Rees M, Sobottka B, et al. A comprehensive single-cell map of T cell exhaustion-associated immune environments in human breast cancer. Nature communications. 2023;14(1):98.

40. Sumida TS, Dulberg S, Schupp JC, Lincoln MR, Stillwell HA, Axisa PP, et al. Type I interferon transcriptional network regulates expression of coinhibitory receptors in human T cells. Nat Immunol. 2022;23(4):632–42.

41. Shao L, Hou W, Scharping NE, Vendetti FP, Srivastava R, Roy CN, et al. IRF1 Inhibits Antitumor Immunity through the Upregulation of PD-L1 in the Tumor Cell. Cancer immunology research. 2019;7(8):1258–66.

42. Man K, Gabriel SS, Liao Y, Gloury R, Preston S, Henstridge DC, et al. Transcription Factor IRF4 Promotes CD8(+) T Cell Exhaustion and Limits the Development of Memory-like T Cells during Chronic Infection. Immunity. 2017;47(6):1129–41 e5.

43. Seo H, Gonzalez-Avalos E, Zhang W, Ramchandani P, Yang C, Lio CJ, et al. BATF and IRF4 cooperate to counter exhaustion in tumor-infiltrating CAR T cells. Nat Immunol. 2021;22(8):983–95.

44. Lynn RC, Weber EW, Sotillo E, Gennert D, Xu P, Good Z, et al. c-Jun overexpression in CAR T cells induces exhaustion resistance. Nature. 2019;576(7786):293-300.

45. Karin M, Liu Z, and Zandi E. AP-1 function and regulation. Curr Opin Cell Biol. 1997;9(2):240–6.

46. Kim YC, and Guan KL. mTOR: a pharmacologic target for autophagy regulation. The Journal of clinical investigation. 2015;125(1):25–32.

47. Yao C, Sun HW, Lacey NE, Ji Y, Moseman EA, Shih HY, et al. Single-cell RNA-seq reveals TOX as a key regulator of CD8(+) T cell persistence in chronic infection. Nat Immunol. 2019;20(7):890–901.

48. Dickinson BC, and Chang CJ. Chemistry and biology of reactive oxygen species in signaling or stress responses. Nat Chem Biol. 2011;7(8):504–11.

49. Finkel T, and Holbrook NJ. Oxidants, oxidative stress and the biology of ageing. Nature. 2000;408(6809):239-47.

50. Wu H, Zhao X, Hochrein SM, Eckstein M, Gubert GF, Knopper K, et al. Mitochondrial dysfunction promotes the transition of precursor to terminally exhausted T cells through HIF-1alpha-mediated glycolytic reprogramming. Nature communications. 2023;14(1):6858.

51. Peng HY, Lucavs J, Ballard D, Das JK, Kumar A, Wang L, et al. Metabolic Reprogramming and Reactive Oxygen Species in T Cell Immunity. Frontiers in immunology. 2021;12:652687.

52. Yu YR, Imrichova H, Wang H, Chao T, Xiao Z, Gao M, et al. Disturbed mitochondrial dynamics in CD8(+) TILs reinforce T cell exhaustion. Nat Immunol. 2020;21(12):1540–51.

53. Scharping NE, Menk AV, Moreci RS, Whetstone RD, Dadey RE, Watkins SC, et al. The Tumor Microenvironment Represses T Cell Mitochondrial Biogenesis to Drive Intratumoral T Cell Metabolic Insufficiency and Dysfunction. Immunity. 2016;45(2):374–88.

54. Bengsch B, Johnson AL, Kurachi M, Odorizzi PM, Pauken KE, Attanasio J, et al. Bioenergetic Insufficiencies Due to Metabolic Alterations Regulated by the Inhibitory Receptor PD-1 Are an Early Driver of CD8(+) T Cell Exhaustion. Immunity. 2016;45(2):358–73.

55. Abeynaike SA, Huynh TR, Mehmood A, Kim T, Frank K, Gao K, et al. Human Hematopoietic Stem Cell Engrafted IL-15 Transgenic NSG Mice Support Robust NK Cell Responses and Sustained HIV-1 Infection. Viruses. 2023;15(2).

56. Aryee KE, Burzenski LM, Yao LC, Keck JG, Greiner DL, Shultz LD, et al. Enhanced development of functional human NK cells in NOD-scid-IL2rg(null) mice expressing human IL15. FASEB journal : official publication of the Federation of American Societies for Experimental Biology. 2022;36(9):e22476.

57. June CH, O’Connor RS, Kawalekar OU, Ghassemi S, and Milone MC. CAR T cell immunotherapy for human cancer. Science. 2018;359(6382):1361–5.

58. Deeks SG, Wagner B, Anton PA, Mitsuyasu RT, Scadden DT, Huang C, et al. A phase II randomized study of HIV-specific T-cell gene therapy in subjects with undetectable plasma viremia on combination antiretroviral therapy. Molecular therapy : the journal of the American Society of Gene Therapy. 2002;5(6):788–97.

59. Mitsuyasu RT, Anton PA, Deeks SG, Scadden DT, Connick E, Downs MT, et al. Prolonged survival and tissue trafficking following adoptive transfer of CD4zeta gene-modified autologous CD4(+) and CD8(+) T cells in human immunodeficiency virus-infected subjects. Blood. 2000;96(3):785–93.

60. Walker RE, Bechtel CM, Natarajan V, Baseler M, Hege KM, Metcalf JA, et al. Long-term in vivo survival of receptor-modified syngeneic T cells in patients with human immunodeficiency virus infection. Blood. 2000;96(2):467–74.

61. Beauparlant D, Rusert P, Magnus C, Kadelka C, Weber J, Uhr T, et al. Delineating CD4 dependency of HIV-1: Adaptation to infect low level CD4 expressing target cells widens cellular tropism but severely impacts on envelope functionality. PLoS Pathog. 2017;13(3):e1006255.

62. Leibman RS, Richardson MW, Ellebrecht CT, Maldini CR, Glover JA, Secreto AJ, et al. Supraphysiologic control over HIV-1 replication mediated by CD8 T cells expressing a re-engineered CD4-based chimeric antigen receptor. PLoS pathogens. 2017;13(10):e1006613.

63. Anthony-Gonda K, Ray A, Su H, Wang Y, Xiong Y, Lee D, et al. In vivo killing of primary HIV-infected cells by peripheral-injected early memory-enriched anti-HIV duoCAR T cells. JCI insight. 2022;7(21).

64. Anthony-Gonda K, Bardhi A, Ray A, Flerin N, Li M, Chen W, et al. Multispecific anti-HIV duoCAR-T cells display broad in vitro antiviral activity and potent in vivo elimination of HIV-infected cells in a humanized mouse model. Science translational medicine. 2019;11(504).

65. Barber-Axthelm IM, Barber-Axthelm V, Sze KY, Zhen A, Suryawanshi GW, Chen IS, et al. Stem cell-derived CAR T cells traffic to HIV reservoirs in macaques. JCI insight. 2021;6(1).

66. Carrillo MA, Zhen A, Zack JA, and Kitchen SG. New approaches for the enhancement of chimeric antigen receptors for the treatment of HIV. Transl Res. 2017;187:83–92.

67. Zhen A, Carrillo MA, and Kitchen SG. Chimeric antigen receptor engineered stem cells: a novel HIV therapy. Immunotherapy. 2017;9(5):401–10.

68. Zhen A, Rezek V, Youn C, Rick J, Lam B, Chang N, et al. Stem-cell Based Engineered Immunity Against HIV Infection in the Humanized Mouse Model. Journal of visualized experiments : JoVE. 2016(113).

69. Mu W, Patankar V, Kitchen S, and Zhen A. Examining Chronic Inflammation, Immune Metabolism, and T Cell Dysfunction in HIV Infection. Viruses. 2024;16(2).

70. Fenwick C, Joo V, Jacquier P, Noto A, Banga R, Perreau M, et al. T-cell exhaustion in HIV infection. Immunological reviews. 2019;292(1):149–63.

71. Zhen A, Rezek V, Youn C, Lam B, Chang N, Rick J, et al. Targeting type I interferon-mediated activation restores immune function in chronic HIV infection. The Journal of clinical investigation. 2017;127(1):260–8.

72. Mu W, Martin H, and Zhen A. Targeting autophagy to treat HIV immune dysfunction. Autophagy Rep. 2023;2(1).

73. Lee DJW, Hodzic Kuerec A, and Maier AB. Targeting ageing with rapamycin and its derivatives in humans: a systematic review. Lancet Healthy Longev. 2024;5(2):e152–e62.

74. Rodes B, Cadinanos J, Esteban-Cantos A, Rodriguez-Centeno J, and Arribas JR. Ageing with HIV: Challenges and biomarkers. EBioMedicine. 2022;77:103896.

75. Babu H, Ambikan AT, Gabriel EE, Svensson Akusjarvi S, Palaniappan AN, Sundaraj V, et al. Systemic Inflammation and the Increased Risk of Inflamm-Aging and Age-Associated Diseases in People Living With HIV on Long Term Suppressive Antiretroviral Therapy. Frontiers in immunology. 2019;10:1965.

76. Nasi M, De Biasi S, Gibellini L, Bianchini E, Pecorini S, Bacca V, et al. Ageing and inflammation in patients with HIV infection. Clin Exp Immunol. 2017;187(1):44–52.

77. Stock PG, Barin B, Hatano H, Rogers RL, Roland ME, Lee TH, et al. Reduction of HIV persistence following transplantation in HIV-infected kidney transplant recipients. Am J Transplant. 2014;14(5):1136–41.

78. Chi H. Regulation and function of mTOR signalling in T cell fate decisions. Nature reviews Immunology. 2012;12(5):325–38.

79. Varco-Merth BD, Brantley W, Marenco A, Duell DD, Fachko DN, Richardson B, et al. Rapamycin limits CD4+ T cell proliferation in simian immunodeficiency virus-infected rhesus macaques on antiretroviral therapy. The Journal of clinical investigation. 2022;132(10).

80. Kouro T, Himuro H, and Sasada T. Exhaustion of CAR T cells: potential causes and solutions. Journal of translational medicine. 2022;20(1):239.

81. Mu W, Zhen A, Carrillo MA, Rezek V, Martin H, Lizarraga M, et al. Oral Combinational Antiretroviral Treatment in HIV-1 Infected Humanized Mice. Journal of visualized experiments : JoVE. 2022(188).

