## supplemental figure for "Rapamycin Enhances CAR-T Control of HIV Replication and Reservoir Elimination *in vivo*"

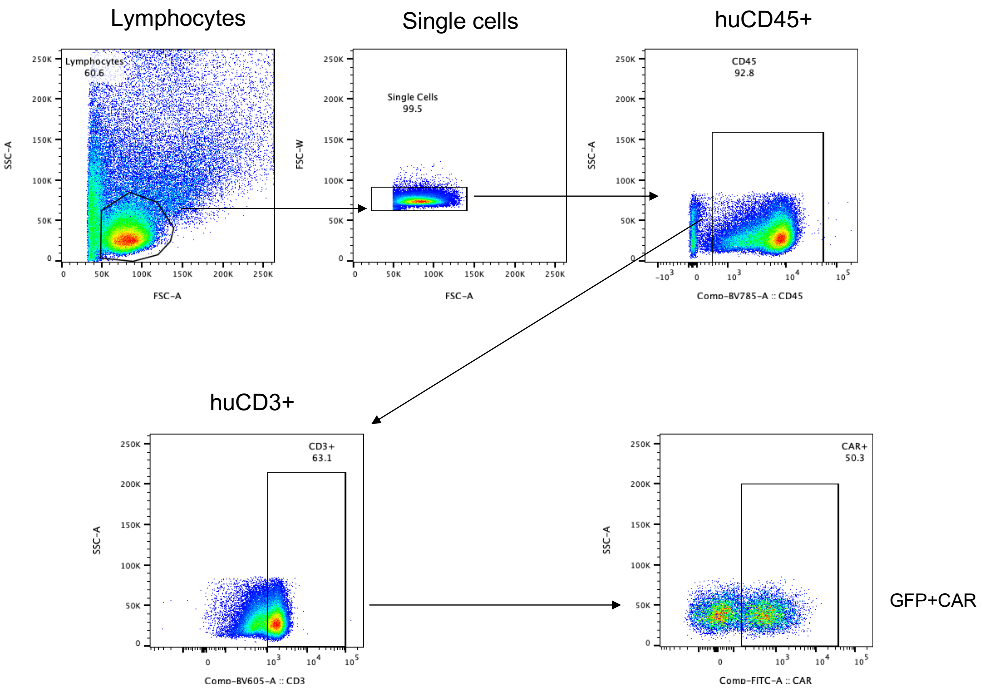


**SFig.1** Representative flow gating huCD45+CD3+GFP+CAR T by flow cytometry from peripheral blood.
